# Season-specific carry-over of early-life associations in a monogamous bird species

**DOI:** 10.1101/785345

**Authors:** Ralf H.J.M. Kurvers, Lea Prox, Damien R. Farine, Coretta Jongeling, Lysanne Snijders

## Abstract

Social relationships can have important fitness consequences. Although there is increasing evidence that social relationships carry over across contexts, few studies have investigated whether relationships formed early in life are carried over to adulthood. For example, juveniles of monogamous species go through a major life-history stage transition—pair formation—during which the pair bond becomes a central unit of the social organization. At present, it remains unclear if pair members retain their early-life relationships after pair formation. We investigated whether same-sex associations formed early in life carry over into adulthood and whether carry-over was dependent on season, in a monogamous species. Moreover, we investigated the role of familiarity, genetic relatedness and aggression on the perseverance of social associations. We studied the social structure before and after pair formation in captive barnacle geese (*Branta leucopsis*), a highly social, long-lived, monogamous species. We constructed association networks of groups of geese before pair formation, during the subsequent breeding season, and in the following wintering season. Next, we studied how these associations carried over during seasonal changes. We found that early-life associations in females were lost during the breeding season, but resurfaced during the subsequent wintering season. In males, the early-life associations persisted across both seasons. Association persistence was not mediated by genetic relatedness or familiarity. The high level of aggressiveness of males, but not females, in the breeding season suggests that males may have played a key role in shaping both their own social environment and that of their partners. We show that early-life social relationships can be maintained well into later life. Such relationships can be sustained even if they are temporarily disrupted, for example due to reproductive behaviour. Our findings therefore highlight that the early-life social environment can have life-long consequences on individuals’ social environment.

## Introduction

The adaptive nature of sociality is a long-standing topic in ecology and evolution (Alexander, 1974; Krause & Ruxton, 2002), particularly the potential costs and benefits of maintaining stable social relationships. Repeated social interactions between the same individuals have, for example, been linked to faster predator-evasion responses, increased foraging success (Carter et al., 2009; Griffiths et al., 2004), and can, ultimately, affect fitness (Beletsky & Orians, 1989; Cameron et al., 2009; Kohn, 2017; Silk, 2007; Silk et al., 2009, 2010).

There is increasing evidence showing that social relationships carry over across time, place, and context. Shizuka et al. (2014) showed that golden-crowned sparrows (*Zonotrichia atricapilla*) that flocked together in one winter flocked together in the subsequent winter more often than expected based on the degree of home range overlap. Roosting associations in Bechstein’s bats (*Myotis bechsteinii*) and Natterer’s bats (*Myotis nattereri*) were found to remain stable across several years despite high fission-fusion dynamics (Kerth et al., 2011; Zeus et al., 2018). Stanley et al. (2018) revealed that, despite seasonal fluctuations in gregariousness and overall weak social associations, semi-feral ponies (*Equus caballus*) maintained stable association preferences over three years. Finally, Firth and Sheldon (2016) showed that great tits’ (*Parus major*) winter social associations carried over into their subsequent breeding season, as individuals bred nearer to those they were most associated with during winter. Despite evidence that social associations between individuals can persist across time and context, it is still largely unknown when these relationships are formed in the first place.

Experiences in early life are known to regularly carry over to adulthood, influencing survival and reproductive performance (Lindström, 1999). Early-life social conditions are generally known to have long-lasting effects (Langenhof & Komdeur, 2018; Leris & Reader, 2016; Stanton & Mann, 2012; Szipl et al., 2019). Despite the likely significance of the early-life period for the formation of social relationships, only a few studies have tracked the social associations of the same individuals from their early life into adulthood (e.g., Carter et al., 2013; Linklater & Cameron, 2009; Mitani, 2009), and almost exclusively in mammals (but see Frigerio et al., 2001 for long-term associations between female siblings in geese). As they mature, individuals of many taxa go through major life-history transitions, such as dispersal (Blumstein et al., 2009; Linklater & Cameron, 2009) and pair formation (Kurvers et al., 2013). These transitions will strongly impact the structure of an individual’s social environment, but perturbations in the social environment do not necessarily impact the stability of already-established relationships (e.g., Kerth et al., 2011). It remains an open question if a life-history transition such as pair formation will cause preferred associations formed in early life to disappear or whether these may persist into adulthood.

Here, we examined the social structure of barnacle geese, a highly social species, before and after pair formation, in order to study the persistence (or disappearance) of early-life same-sex associations after a major life-history transition: pair formation. Barnacle geese are long-lived monogamous birds, which generally find a partner at 2–3 years of age (Choudhury & Black, 1994). Barnacle geese are very selective in choosing a mate; they sample one to six potential mates in so-called “trial liaisons” before settling with a permanent partner (Choudhury & Black, 1993; van der Jeugd & Blaakmeer, 2001). The pair bond is extremely strong and pair members generally stay in close proximity to each other and remain together until one of them dies (Black, 2001; Owen et al., 1988). Pairs with a longer pair bond show higher reproductive success than pairs with a shorter pair bond (Black, 2001). Pair formation is thus a crucial step in the life history of geese. Here we took advantage of controlled experiments to study how pair formation impacted same-sex foraging associations formed early in life, allowing us to circumvent problems with missing individuals in naturalistic data. We followed a captive population of barnacle geese over several years, quantifying the dyadic association strength between individual geese before and after pair formation. To study mediating factors underlying the potential persistence of these dyadic associations, we furthermore quantified the degree of familiarity and genetic relatedness in the population, as well as individuals’ aggressive interactions. We focused on the persistence of same-sex associations because, in comparison to inter-sex associations, these relationships are less well studied in monogamous social species, but may have important adaptive benefits, for example stress-reduction (Sachser et al., 1998), especially in competitive contexts such as foraging.

In the wild, barnacle geese, like many migratory waterfowl, show a high site fidelity to their breeding, wintering and staging areas (Percival, 1991; Robertson & Cooke, 1999), creating opportunities for long-term maintenance of social relationships established early in life. Both sexes show natal philopatry, though females have substantially higher levels of natal philopatry than males (van der Jeugd, 2001; van der Jeugd et al., 2002). The extent to which individuals maintain non-sexual relationships across seasons is not well understood, partly because of the challenges of following individuals across space and time. van der Jeugd et al. (2002) showed that during breeding, female, but not male, barnacle geese nested closer to siblings than expected by chance. This occurred not only when the female siblings bred in the same island as their parents, but also when they nested on a different island. Moreover, this relationship was only observed for siblings hatched in the same year. Similar sex-specific patterns have been observed in terms of resting proximity in semi-feral greylag geese, *Anser anser*, in winter (Frigerio et al., 2001). Given these observations and females being the more philopatric sex, we predicted that females would show a higher likelihood of maintaining their early-life relationships than males after mating.

We constructed association matrices from foraging observations collected during the breeding and wintering seasons. Geese are generally more gregarious during the wintering season (Black et al., 2014; Szipl et al., 2019), while being more aggressive and territorial during the breeding season (Owen & Wells, 1979). We, therefore, predicted that associations with early-life companions after pair formation would be stronger in the wintering than the breeding season. Finally, based on the expected direct and indirect adaptive benefits of associating with kin, for example by providing and receiving social support (Black & Owen, 1989; Raveling et al., 2000; Scheiber et al., 2009; Scheiber et al., 2005), we predicted that genetic relatedness would positively impact the strength of the associations. Similarly, based on the expected benefits of reduced aggression between familiar individuals (i.e. “dear enemy effect”; van der Jeugd, 2001; Ydenberg et al., 1988), we predicted that long-term familiarity would positively influence dyadic foraging association strength.

## Methods

### Study subjects

In late 2007 we obtained two mixed-sex groups of barnacle geese (see Fig. 1 for study timeline, and Kurvers et al., 2013 for more details). The two groups consisted of 21 (13 female, 8 male) and 23 individuals (8 female, 15 male) respectively. The groups were housed in separate outdoor aviaries (12 × 15 m) at the Netherlands Institute of Ecology (NIOO) in Heteren, the Netherlands. Groups were visually, but not acoustically, isolated from each other. The aviaries consisted of bare soil with a large pond (6 × 1 m) with running water for drinking and bathing. Birds received ad libitum food consisting of a mixture of grains and pellets occasionally supplemented with grass. Most individuals (40 of 44) were hatched in 2007 and were thus approximately 5 months old upon arrival. All birds were captive-hatched, wing-clipped and (upon arrival) fitted with uniquely coded white leg rings for identification. Birds from the two groups had different origins, implying that birds within (but not between) groups could have a high genetic relatedness (see below). Birds within a group will henceforth be referred to as “familiar” individuals, and birds between groups as “unfamiliar”. Geese lived for approximately 1.5 years in these familiarity groups before the start of the social network observations (see below).

**Figure 1.**
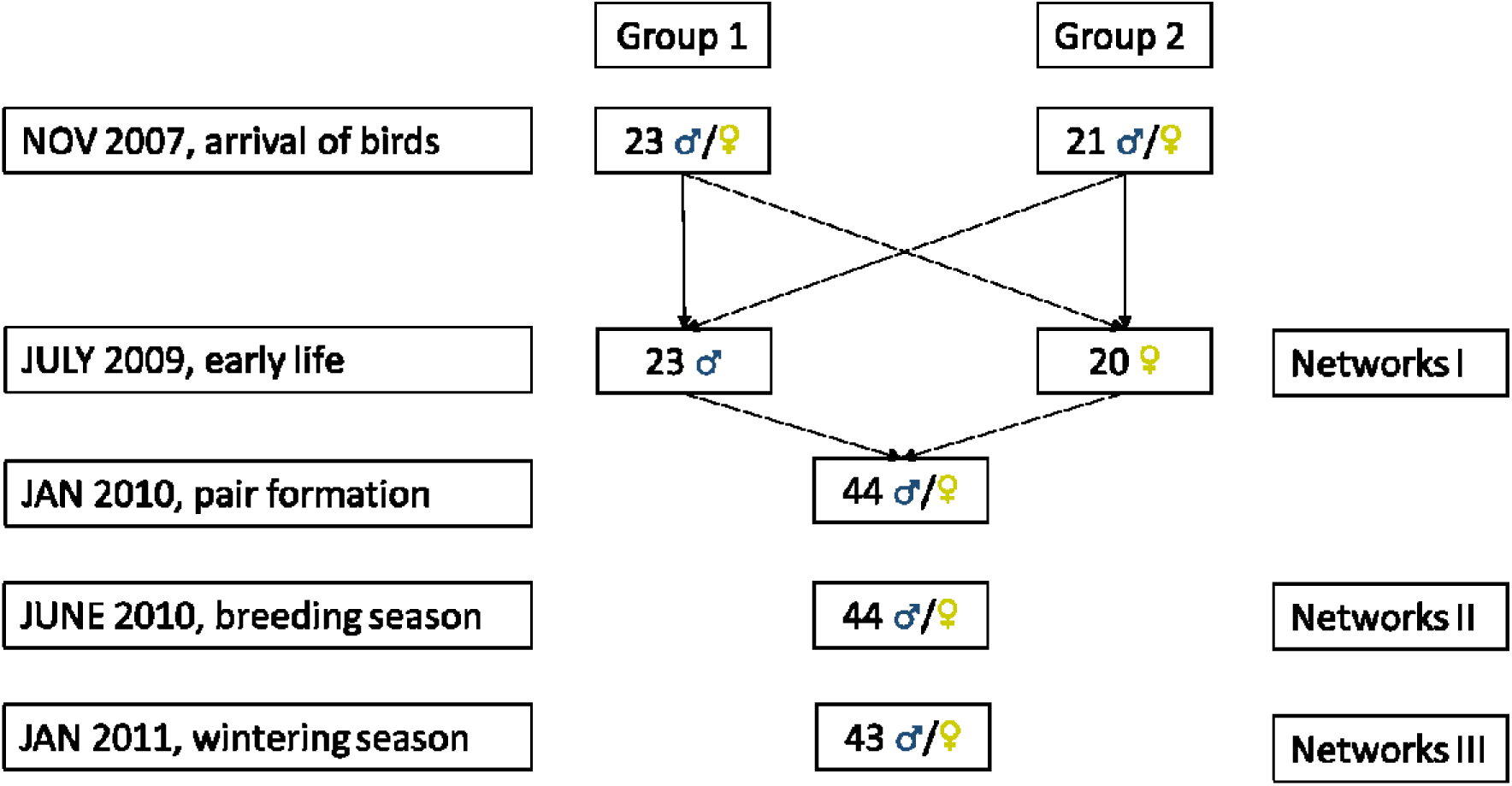
Schematic overview of the experimental procedures. Geese were procured in November 2007—at which point almost all were juveniles (hatched in 2007)—and placed in one of two familiarity groups. In July 2009, males and females of both groups were placed together and we quantified the social network structure of both groups (to examine the role of familiarity and genetic relatedness on social structure, Kurvers et al., 2013). In January 2010, all geese were placed together in one group and individuals rapidly started to form pairs. In the subsequent breeding season (June 2010) and wintering season (January 2011), we quantified the social network structure of the entire group. Note that the sex of one individual could not be reliably determined at the time of the first network observation and hence was not included in the early-life observations. Another individual died in between the breeding and wintering season. Both individuals were excluded from all analyses, resulting in a final sample size of 42 (20 females, 22 males).

### Genetic relatedness

We determined the genetic relatedness between each pair of individuals using a high-resolution 374 single nucleotide polymorphism (SNP) marker set, developed for the barnacle goose (Jonker et al., 2012). We took a small blood sample (± 1 ml) from each individual from the brachial vein and preserved it in ethanol. Entire genomic DNA was isolated using the Gentra Systems Puregene DNA purification kit. Genotyping was conducted with Vera Code assays on an Illumina BeadXpress (described in Kraus et al., 2011). We calculated the pairwise genetic relatedness (*r*) using the program Coancestry (version 1, Wang, 2011). To determine which relatedness estimator best fitted our data, we used the empirical SNP allele frequencies of our population and simulated 500 dyads of geese of varying relatedness coefficients. Meeting real conditions as closely as possible, we simulated mainly unrelated dyads (*N* = 430) but also dyads of close familial relationships, namely 30 full sibs dyads (*r =* 0.5), 10 half sibs dyads (*r =* 0.25), and 30 first cousins dyads (*r =* 0.125). Based on this simulation, we found that the maximum-likelihood estimator of Milligan (2003) performed best and used it for the final estimation of *r*. This produced a strong correlation with expected values of *r* (*r*^2^ = 0.9; analysis of the simulated data set carried out with default settings). Subsequently, all pairwise relatedness values of the experimental geese were obtained from Coancestry with standard settings (see Kurvers et al., 2013 for more details).

### Observations prior to pair formation: early life

After living for approximately 1.5 years in their respective familiarity groups—and prior to any pair formation—we separated all geese into two single-sex groups (June 2009; Fig. 1) to study the factors shaping intra-sexual social association preferences while avoiding additional ‘noise’ caused by inter-sexual trial-liaisons (van der Jeugd & Blaakmeer, 2001). Moreover, by separating individuals in single-sex groups, we could control the timing of pair formation and keep this comparable and tractable for all subjects. Geese were sexed by visual inspection of sexual organs in the cloaca. The sex of one individual could not be reliably determined at this time. This individual was not included in the early-life observations, but joined the flock after these observations finished. This individual was excluded from all analyses. Association observations were conducted in the home aviaries during foraging. These foraging associations were studied on five grass patches (40 × 20 cm, 1.5m apart) which were replaced twice a day to avoid depletion. Other food sources were removed during the observations.

Each single-sex group was observed (9 a.m.–1 p.m.) for 15 days (females: 22 June–12 July 2009; males: 13–30 July 2009). The presence of all individuals on the patches was recorded every 4 minutes. This interval was longer than the mixing time among individuals (i.e., the time individuals need to exchange who they are association with), in order to ensure independence of observations (Croft et al., 2008). Associations at feeding patches was rarely the same in consecutive records (females: 5.9%; males: 7.9%). Observations were occasionally interrupted for 10 minutes in the event of an external disturbance. Since patch size (40 × 20 cm) and group size (mean females = 1.9, range 1–5; mean males = 2.0, range 1– 5; Fig. A1a, b) were small, we assumed that animals grazing on the same patch during a sampling period were associating (a.k.a. gambit of the group, Franks et al., 2010; Whitehead, 2008).

The results of the foraging associations before pair formation are reported in Kurvers et al. (2013). In brief, in both sexes, familiarity and genetic relatedness predicted association strength, whereas boldness and dominance did not. Therefore, we focus here on the role of familiarity and genetic relatedness in governing the stability of long-term associations.

### Observations after pair formation: breeding and wintering

Geese remained in the single-sex groups for six months and were then placed together in one group (January 2010; Fig. 1). By then, most geese were 2.5 years of age, approximating the average age of final pair formation in barnacle geese (Choudhury & Black, 1994; van der Jeugd & Blaakmeer, 2001). Pair formation started rapidly, and most geese quickly formed a stable pair bond. Before starting the first post-pair formation association observations, 37 (of 44) geese had formed stable pair bonds. In total, geese formed 12 pairs (11 male-female pairs, and one male-male pair), four triplets (three geese continuously moving together as one unit without any aggression; two triplets consisted of two females and one male, and two triplets of one female and two males) and seven geese (3 males, 4 females) remained unpaired. The occurrence of triplets has also been observed in the wild, with a third party joining a pair between 10 months and 4 years, however incidence in the field was lower (Black et al., 1996). As reported in Kurvers et al. (2013) genetic relatedness did not play a role in mate choice, whereas geese actively selected against familiarity in selecting a mate. After the geese formed pair relationships, and during their first breeding season, we again conducted observations of foraging associations. We placed 10 grass patches (40 × 40 cm) in the aviary. Observations were conducted for five weeks (31 May–2 July 2010, 25 observation days) in two 2-hour blocks per day (8 a.m.–1 p.m.) following the same observation protocol described above. During the breeding season, many of the paired individuals started building nests and laying eggs. We regularly checked all nests and removed any eggs to avoid the hatching of goslings.

To study the stability of associations across seasons, we repeated the observations of foraging associations six months later, during the wintering season, following the same protocol (17 December 2010–4 February 2011, 13 observation days). Comparing the pair status of all geese between the breeding and wintering seasons, we observed that all pair relationships but one remained the same—the one pair relationship that changed was the male-male pair. This reflects the strong and long lasting pair bonds in barnacle geese, which generally stay together until one of the pair members dies (Black, 2001). One individual died in between the breeding and wintering observations. To facilitate comparison across seasons, we removed this individual from all analyses. The final sample size was thus 42 individuals (20 females, 22 males).

### Agonistic interactions

During all three observation periods (early life, breeding, wintering), we collected data on agonistic interactions to study the role of aggression on the stability of early-life associations. In between scoring the presence of individuals on the patches, we scored the winner and loser of agonistic interactions, defined as a direct confrontation between two geese and ranging from threats with lowered head and neck to active chases with flapping wings. In the early-life observations, we identified a total of 1,429 interactions in the female group and 2,619 interactions in the male group. We then identified 3,411 and 786 interactions during the subsequent breeding and wintering periods, respectively.

### Statistical analyses

#### Stability of early-life connections

We used social network analysis to investigate carry-over effects of dyadic associations between seasons. Within the networks, nodes represent individuals and are connected by edges that represent associations. The edge weight varied with dyadic association strength. From the observation data, we thus generated undirected weighted networks (i.e., networks based on associations without initiators or receivers) for each of the three periods (early life, breeding season, and wintering season). Edge weights in the networks were calculated using the simple ratio index (SRI) as an association measure using the ‘asnipe’ package (Farine, 2013) in R (v. 3.4.4). The SRI indicates the probability of observing two individuals in association with each other given that one was observed. Values range from 0 (two individuals were never observed together) to 1 (two individuals were always observed together). The SRI is considered an effective measure of dyadic association strength provided there are no large sampling biases (Farine & Whitehead, 2015; Ginsberg & Young, 1992; Hoppitt & Farine, 2018). Since all observations were performed within aviaries in which all individuals feeding on all patches could be easily observed, we did not expect a strong sampling bias.

We first analysed whether early-life associations (within the same sex) were maintained after pair formation, i.e. during the subsequent breeding and wintering season. Within each sex, we performed Mantel tests using the built-in node permutation test from the ‘vegan’ library Mantel test function—running 50,000 permutations and using the spearman correlation (Oksanen et al., 2019)—to compare dyads’ association strength between (i) early life and breeding season, and (ii) early life and wintering season. We started with a separate matrix for each sex for the early-life associations, one matrix (containing both sexes) for the breeding season, and one matrix for the wintering season. To compare the matrix correlation for females (/males), we therefore first subset all females (/males) from the breeding and wintering matrix, constructing single-sex matrices. To confirm that our results were not mainly driven by the associations of the unpaired individuals (as we were primarily interested in the possible continuation of associations after pair formation), we repeated these analyses after further excluding associations between the unpaired individuals. To investigate if our results were driven by a few strong associations, we also repeated this analysis while excluding all associations with an SRI value higher than 0.1.

#### Genetic relatedness and familiarity

To investigate which factors might underlie a possible continuation of associations after pair formation, we studied the effect of genetic relatedness and familiarity on dyadic association strength during the breeding and wintering season. We tested the effect of familiarity and relatedness using multiple regression quadratic assignment procedures (MRQAP; Krackhardt, 1988) using the mrqap.dsp function in the R package asnipe (Farine, 2013). With this procedure, the association matrix was first regressed against two matrices, one with data on relatedness and one on familiarity. We used a version of the function allowing us to combine the mrqap.dsp model with pre-network permutations (Farine, 2017). Specifically, we generated 1,000 random networks by re-organising the observations of individuals in the original group-by-individual matrices following the method first described by Bejder et al. (1998). We then compared the resulting distribution of coefficient values from these permuted networks to the coefficient value generated from the original observation data to obtain *P*-values. Further, because our random distributions were not centred on 0 (see Farine, 2017), we rescaled measures to an effect size by taking the difference between the observed coefficient values and the mean of the corresponding distribution of coefficient values based on the permutated networks. We performed the MRQAP separately for each season and sex.

#### Patch visits and aggression

As we found sex differences in the continuation of early-life associations (see below), we investigated the role of patch visitation rates and aggression as potential underlying mechanisms. For each individual, for each period, we calculated the mean number of patch visits per hour. We then used a Mann-Whitney U test to determine whether the sexes differed in their patch visitation rates in each of the three seasons. Similarly, for each individual and for each season, we determined the mean number of aggressive interactions initiated per hour, and tested whether the sexes differed in their likelihood to display aggression in each of the three seasons.

### Ethical permission

The animal ethical committee of both the Royal Netherlands Academy of Arts and Sciences (KNAW) and the Wageningen University approved all experiments [protocol numbers: 2010008.b (blood sampling)]. Geese were obtained from a waterfowl breeding farm (Kooy and Sons, ‘t Zand, the Netherlands) and returned there after the experiments had finished.

## Results

### Sex- and season-dependent effects of persistence of early-life associations

We quantified the foraging association networks for each of the three observation periods (i.e., early life, breeding season, and wintering season, see Fig. 2), and compared the strength of the dyadic associations that were formed early in life to those in subsequent seasons. We found that a female’s dyadic association strengths from early life (i.e. prior to pair formation) were not significantly correlated with its dyadic association strengths to the same pool of individuals (i.e. other females) in the subsequent breeding season (*r =* 0.04; *P =* 0.32; Fig. 3a). A female’s early-life dyadic association strengths were, however, significantly correlated with its dyadic association strengths in the subsequent wintering season (*r =* 0.18; *P =* 0.019; Fig. 3b). A male’s early-life dyadic association strengths were significantly correlated with its dyadic association strengths to other males in both the subsequent breeding (*r =* 0.26; *P =* 0.002; Fig. 3c) and the following wintering season (*r =* 0.31; *P =* 0.002; Fig. 3d).

**Figure 2.**
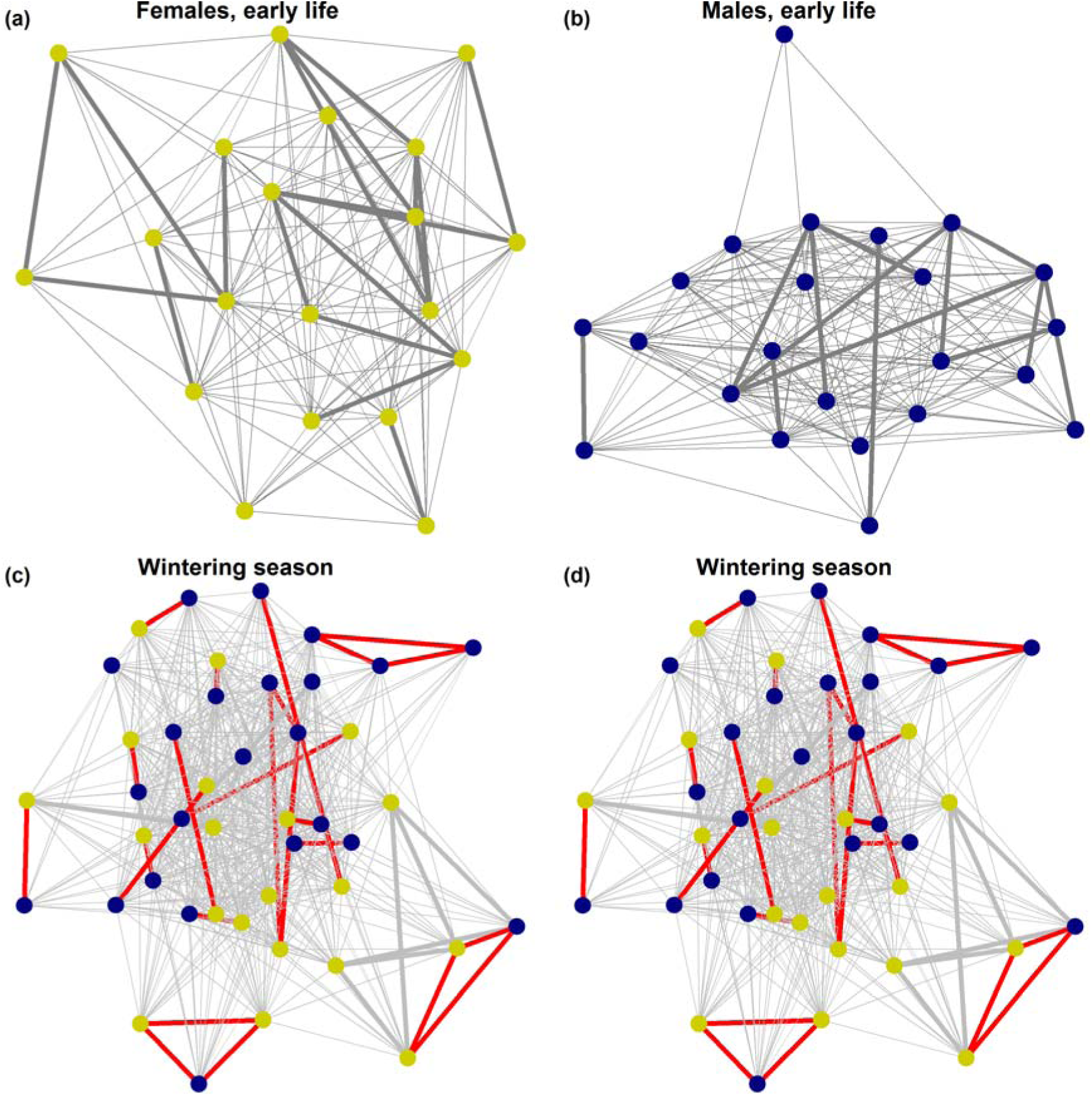
The (a) female and (b) male network in single-sex groups prior to pair formation and the network of all individuals after pair formation during the (c) breeding, and (d) wintering season. Yellow/blue circles represent females/males respectively. For visualization, we removed associations below SRI values of 0.005. Thin/thick lines represent SRI values below/above 0.1. Coloured lines indicate a pair bond. Networks were created with the ggnet2 function in R using the Fruchterman-Reingold algorithm for node placement.

**Figure 3.**
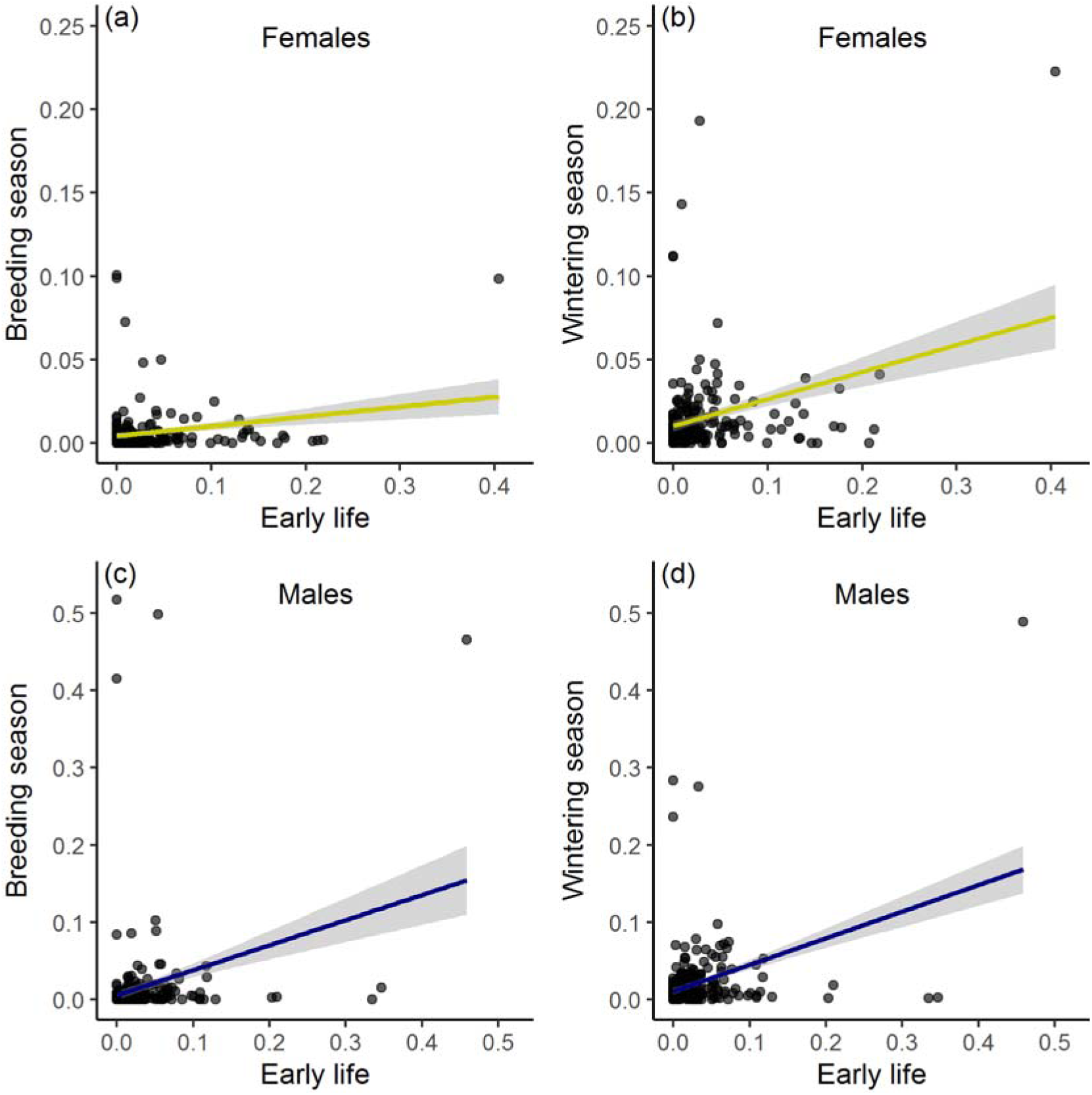
The relationship between the association strength of female dyads comparing (a) early life and breeding season, and (b) early life and wintering season, and the relationship between the association strength of male dyads comparing (c) early life and breeding season, and (d) early life and wintering season. Lines are linear regression lines, including 95% confidence bands.

When excluding the associations among individuals that remained unpaired (7 of 42; but maintaining the associations between paired and unpaired individuals), we obtained similar results (females: early life vs. breeding: *r =* 0.00, *P =* 0.49; early life vs. wintering: *r =* 0.19, *P =* 0.018; males: early life vs. breeding: *r =* 0.26, *P =* 0.003; early life vs. wintering: *r =* 0.31, *P =* 0.003).

When excluding all dyadic associations with an SRI value above 0.1 (which excluded 38 associations for females (7% of all associations) and 26 for males (5%)), we, again, obtained similar results (females: early life vs. breeding: *r =* 0.00, *P =* 0.52; early life vs. wintering: *r =* 0.18, *P =* 0.022; males: early life vs. breeding: *r =* 0.23, *P =* 0.004; early life vs. wintering: *r =* 0.34, *P =* 0.002).

### Genetic relatedness and familiarity do not drive associations in females or males

In females, there was no effect of genetic relatedness or familiarity on dyadic association strength in the breeding season (effect sizes: genetic relatedness: −0.00006; familiarity: 0.0001) or in the wintering season (genetic relatedness: 0.0016; familiarity: 0.0001). Likewise, for males, we found no effect of genetic relatedness or familiarity on dyadic association strength in the breeding season (genetic relatedness: −0.00035; familiarity: - 0.00006) or in the wintering season (genetic relatedness: 0.0002; familiarity: −0.00026). Thus, although familiarity and genetic relatedness positively impacted single-sex associations in females and males in early life (see Kurvers et al., 2013 and see Fig. A2 and Fig. A3), these factors did not drive same-sex associations after pair formation (all *P >* 0.1).

### Elevated aggression in males and during the breeding season

We investigated two mechanisms that might drive the disappearance of social associations in females—but not males—during the breeding season. First, females may simply visit the food patches less than males during the breeding season, which would lower their opportunities for maintaining relationships. Though females visited patches less than males in the single-sex groups before pair-formation (*W* = 50, *P <* 0.001; Fig. 4a), females visited the patches equally often as males in the breeding and wintering season (both *P >* 0.35; Fig. 4b, c), ruling out this explanation.

**Figure 4.**
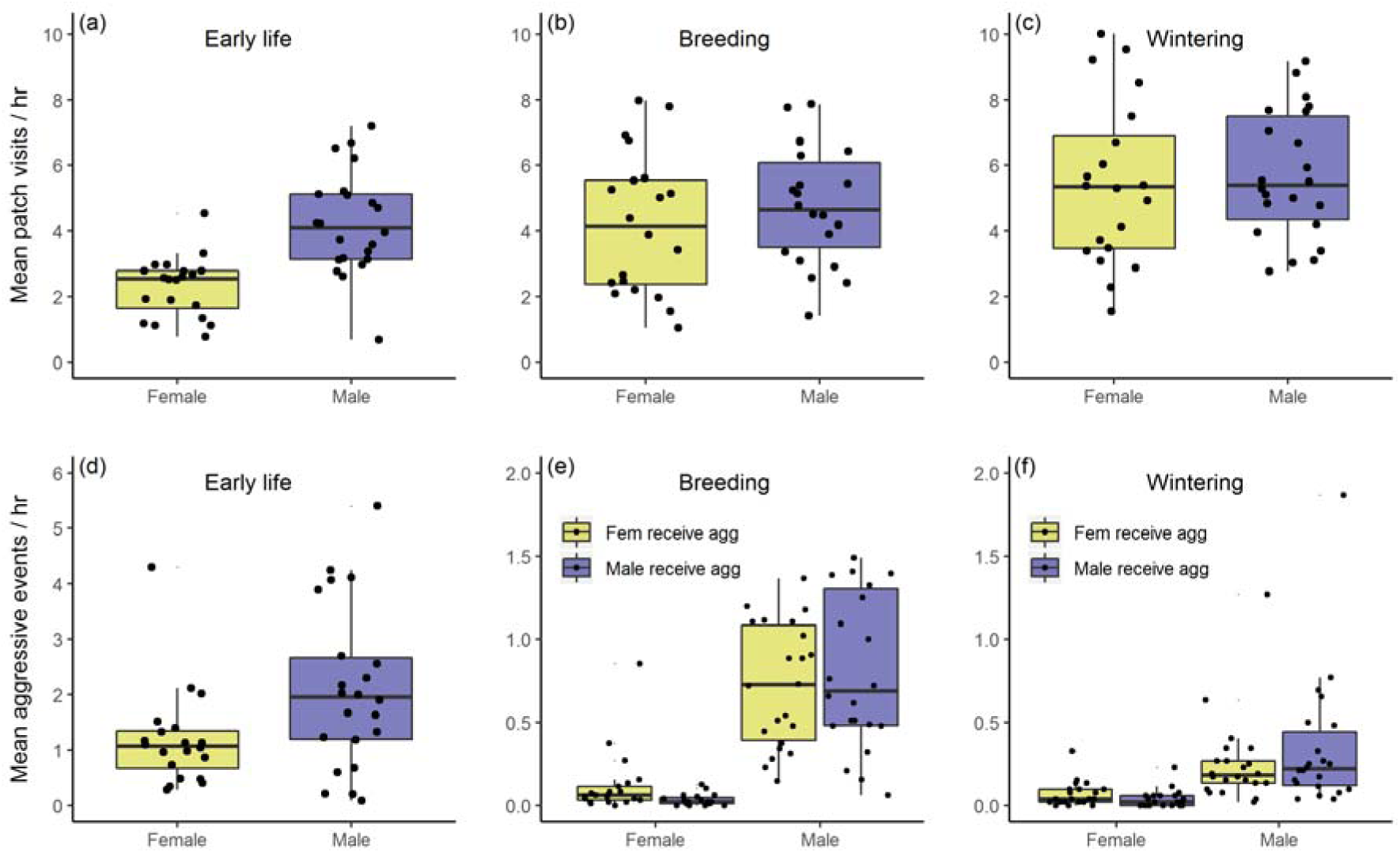
The mean number of patches visited per hour for (a) females and males in the single-sex groups prior to pair formation, and for both sexes during the (b) breeding and (c) wintering seasons after pair formation. The mean number of aggressive events initiated per hour for (d) females and males in the single-sex groups prior to pair formation, and for both sexes during the (e) breeding and (f) wintering season after pair formation. (e, f) Colours indicate the sex of the individual receiving the aggression. Dots represent individuals. Boxplots show median, interquartile ranges and whiskers show the lowest/highest values within the 1.5 interquartile range.

A second mechanism might be that males, being generally the more dominant member of a pair, play a stronger role in determining the association members of a pair than do females, especially in the breeding season. Before pair formation, males showed a higher level of aggression than females in the single-sex groups (*W* = 142, *P =* 0.049; Fig. 4d). Likewise, males showed a substantially higher level of aggression than females during the breeding (*W* = 8, *P <* 0.001; Fig. 4e) and wintering seasons (*W* = 39, *P <* 0.001; Fig. 4f). Males displayed equal levels of aggression towards males and females in the breeding season (Wilcoxon signed-rank test: *V* = 76, *P =* 0.10, Fig. 4e) but slightly higher levels of aggression towards males than females in the wintering season (*V* = 50, *P =* 0.02; Fig. 4f). As expected, male geese displayed higher levels of aggression in the breeding than in the wintering season (Wilcoxon signed-rank test, males only: *V* = 240, *P <* 0.001; Fig. 4e, f). In accordance, the mean group size at patches was higher during the wintering than during the breeding season (mean group size wintering: 2.4; breeding: 2.0; Fig. A1c, d). Moreover, paired individuals were more tolerant to the presence of other individuals (i.e., not belonging to the pair) at a patch during wintering season. In the breeding season, in 80% of cases when pair members were observed together on a patch, there were no other individuals present. In the wintering season, this percentage dropped to 60% (mean number of non-pair individuals at a patch with a pair during breeding: 0.26, during wintering: 0.76; Fig. A4c, d). This is also apparent in the network graphs showing more edges during the wintering season (Fig. 2c, d).

## Discussion

Maintaining stable social associations across time and contexts can have adaptive benefits (Kohn, 2017). Yet the importance of the early-life period for the formation of such long-term stable relationships has so far received little attention. Here we show that early-life same-sex foraging associations can persist after a major life-history transition—pair formation—in a monogamous and long-lived bird species. However, which associations were maintained depended on sex and season. Early-life associations in females were lost during the breeding season, but resurfaced again during the subsequent wintering season. In males, the early-life associations persisted across both seasons. We found no evidence of genetic relatedness or familiarity on association persistence. Elevated male aggression likely influenced the limited number of contacts outside of the pair bond during the breeding season—and thereby the extent to which early-life associations could be maintained, especially by females during the breeding season. Our findings extend the understanding of how social relationships develop and are maintained over different life-history phases and how their importance to individuals may vary with season.

Across taxa, females are well-known for maintaining long-term social relationships (Cameron et al., 2009; Carter et al., 2013; Ilany et al., 2015; Silk et al., 2003), which is commonly attributed to their reproductive strategies. In our study, males, not females, maintained their associations throughout both the breeding and wintering season. Benefits of social relationships in males have been observed in other species: male red-winged blackbirds (*Agelaius phoeniceus*) that bred close to familiar neighbours fledged more offspring (Beletsky & Orians, 1989) and male Assamese macaques (*Macaca assamensis*) with stronger social associations sired more offspring (Schulke et al., 2010). Males can thus clearly form, and benefit from, long-lasting social relationships.

Social relationships in males often take the form of coalitions or alliances (Connor et al., 2017; Gilby et al., 2013; Schulke et al., 2010), benefiting individuals by providing cooperation partners in agonistic interactions with conspecifics. But reduced aggression, for instance, via dear-enemy effects (Temeles, 1994), can likewise be an important benefit of maintaining long-term social relationships, especially in territorial animals (Chuang et al., 2017; Jaeger, 1981; Siracusa et al., 2019). Additionally, repeated association with certain individuals can influence vigilance behaviour, with individuals showing reduced vigilance in the proximity of well-known conspecifics (Carter et al., 2009; Kutsukake, 2006). In a monogamous prey species such as barnacle geese, in which the male spends much of its time during the breeding season on vigilance at the expense of foraging (Forslund, 1993), such benefits can be especially substantial. The expected benefits of maintaining long-term social relationships vary with ecological and social conditions (Connor et al., 2017; Kappeler et al., 2013; Maher & Burger, 2011), and studying the social structure of both sexes in taxa with distinct space use (e.g., natal philopatry), life history (e.g., long life-span), social organization (e.g., fission-fusion), and mating system (e.g., monogamy) characteristics, offers us greater insight into the drivers and constraints of maintaining long-term stable associations in animal societies.

The apparent lack of social association persistence for females in the breeding season is surprising, but adds to our understanding of social flexibility by showing that the persistence of social relationships not only varies between species and individuals (Kappeler et al., 2013) but can also have sex-specific effects across seasons. Female chacma baboons (*Papio hamadryas ursinus*) were similarly found to vary in their social preferences depending on ecological context (Henzi et al., 2009): social preferences were more pronounced when food resources were scarce, but females acted more “gregariously” (i.e., without social preferences) when resources were plentiful. For geese, winter is a time when resources become scarce; barnacle geese, however, are highly gregarious in winter (Black et al., 2014), possibly to access social information on foraging opportunities (Drent & Swierstra, 1977; Kurvers et al., 2009; Kurvers et al., 2010) (but see Kurvers et al., 2014). Alternatively, the fluctuations in female social preferences may have been the result of the male partner unselectively excluding his partner’s—but not his own—early-life companions during the breeding season. The high level of aggression displayed by the males during the breeding season compared to the wintering season supports this hypothesis.

The absence of continued association with early-life companions in females in the breeding season appears to be in contrast to previous findings. An earlier study on wild barnacle geese found that female, but not male, geese exhibited social preferences in terms of nesting proximity, with females nesting closer to familiar females (van der Jeugd et al., 2002). It is possible that nest choice offers female geese an alternative way to maintain early-life social associations, circumventing the potentially controlling influence of their partner during foraging. Possibly, breeding in proximity to familiar and/or related conspecifics provides context-specific benefits that foraging together does not. Intraspecific brood parasitism and adoption is common in waterfowl (Anderholm et al., 2009a; Andersson et al., 2019; Choudhury et al., 1993; Forslund & Larsson, 1995) and breeding close to related individuals may decrease the costs, through inclusive fitness benefits, of having to care for additional offspring. Moreover, breeding in proximity could enable siblings to defend each other’s nests against unrelated brood parasites. Female siblings may also actively or passively support each other in the acquisition of high-quality nest locations. Indeed, in geese, social support from family members is known to give individuals an advantage in competitive interactions, starting from an early age (Black & Owen, 1989; Raveling et al., 2000; Scheiber et al., 2009; Scheiber et al., 2005). Lastly, neighbouring barnacle geese pairs are known to defend their nests together (Black & Owen, 1995) and nesting closely to familiars may facilitate cooperative nest defence (Grabowska-Zhang et al., 2012; Olendorf et al., 2004). Nest predation is a major threat for geese (Drent & Prop, 2008) that exposes the females in particular to considerable predation risk (Samelius & Alisauskas, 2006).

Our captive study design had several important limitations compared to natural settings. First, the group size under study was relatively small compared to natural groups. In natural groups, the number of genetically related and/or familiar individuals may be substantially higher, allowing geese more opportunities to associate with these types of individuals. Second, the space available to our subjects was reduced as compared to natural conditions. This may have led to higher levels of aggression, especially among males, and/or, more and stronger associations as compared to more natural settings. Third, we removed the eggs of breeding females to avoid the undesired hatching of more experimental animals. This may have caused heightened aggression, and may have had repercussions for social relationships. The pair bonds did, however, almost all remain intact till the next wintering season, suggesting that egg removal did not cause major disruption of pair bonds.

Taken together, our findings suggest that different types of social associations may be beneficial in different contexts and that the early-life period can be a crucial time for the formation of these associations. The next step is to disentangle whether individuals actively choose to (re)associate with earlier companions depending on season- and context-dependent benefits, or whether the observed fluctuations in social association persistence are an emergent property following relatively simple season-dependent social processes, such as heightened aggression. The first scenario may have important implications for our understanding of the cognitive abilities of animals. Notably, Scheiber et al. (2011) found that six-week-old juvenile greylag geese can already discriminate between two of their siblings, showing that individual-level recognition is already present from an early age. Our findings here suggest that geese may be able to keep track of multiple types of relationships in a large fission–fusion society, despite extended breaks, supporting similar observations in wild barnacle geese (Black & Owen, 1995), and that they can re-evaluate the benefits of these relationships depending on context. Given the strong evidence for birthplace-independent long-term kin discrimination in both migrating and captive barnacle geese (Anderholm et al., 2009b; Kurvers et al., 2013; van der Jeugd et al., 2002), this level of cognitive ability is certainly feasible and makes the long-lived barnacle goose an interesting study system to further examine such mechanisms. Complex social patterns can be driven by cognitive ability, but also emerge from relatively simple processes (Kappeler, 2019), such as site fidelity combined with season- and sex-dependent aggression. Unravelling how these mechanisms underlie social complexity in a diversity of social systems will be central to our understanding of the evolution of animal societies.

## Acknowledgements

We thank Bart van Lith from the Netherlands Institute of Ecology for the caretaking of the birds, and Chantal Althuizen and Jan Baar for help during the social network observations. We thank Herbert Prins, Ron Ydenberg and Sip van Wieren for their guidance on the experimental design. We thank the Faunafonds and the Koninklijke Nederlands Jagers Vereniging (KNJV) for financial support, and Bart Nolet and Marcel Klaassen for facilitating the animal holding facilities. We thank Rudy Jonker for blood sampling; Henk van der Jeugd for sexing the geese; and Robert Kraus, Richard Crooijmans, Martien Groenen, Qiong Zhang and Bert Dibbits for the genetic lab work. Finally, we thank Deborah Ain for helpful suggestions on the manuscript. D.R.F. was funded by the Max Planck Society and the DFG Centre of Excellence 2117 “Centre for the Advanced Study of Collective Behaviour” (ID: 422037984). L.S. was funded by a postdoc fellowship of the Alexander von Humboldt Stiftung (Ref 3.3 - NLD - 1192631 - HFST-P).

## Data Availability Statement

The raw social association observations, the matrices containing the SRI values per sex per season, the matrices containing the familiarity and genetic relatedness data per sex, and the raw aggression data are uploaded on the Open Science Framework for referee inspection (<u>link</u>). These files will be made public in case of acceptance.

---

**Figure A1.**
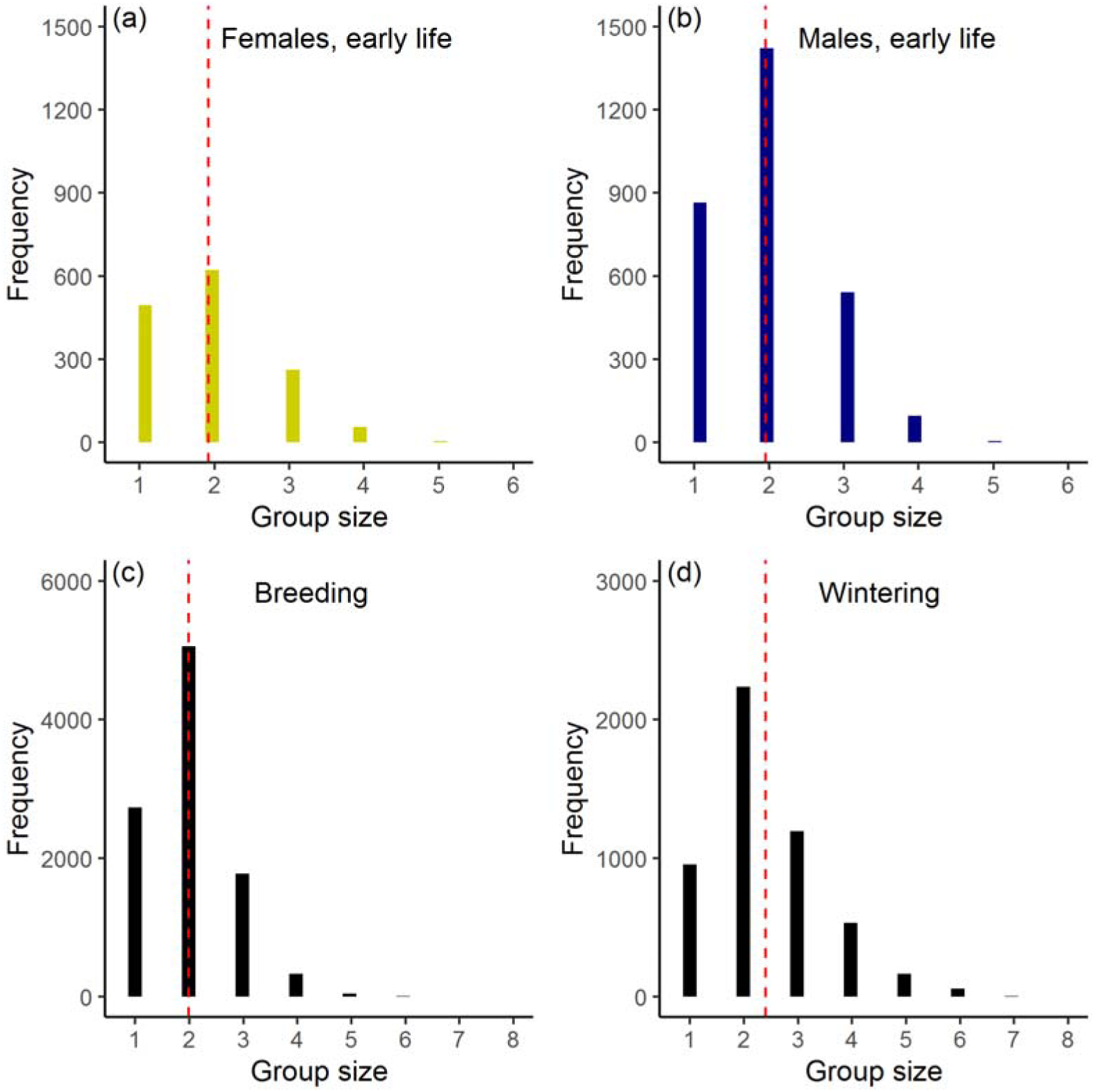
(a, b) The frequency of observed group sizes during the early-life social network observations for (a) females and (b) males. Red dashed lines represent mean group size (mean group size females: 1.92; males: 1.96). Males frequented patches at a higher rate (see also Fig. 4a). (c, d) The frequency of observed group sizes during the (c) breeding and (d) wintering season. Red dashed lines represent mean group size (mean group size breeding: 2.0; wintering: 2.4). Note that the overall higher frequency of group sizes in the breeding season is due to more observation days during breeding (*N* = 25 days) than during wintering season (*N* = 13 days). The mean patch visit rate was in fact slightly higher in wintering than in breeding season (see Fig. 4b, c).

**Figure A2.**
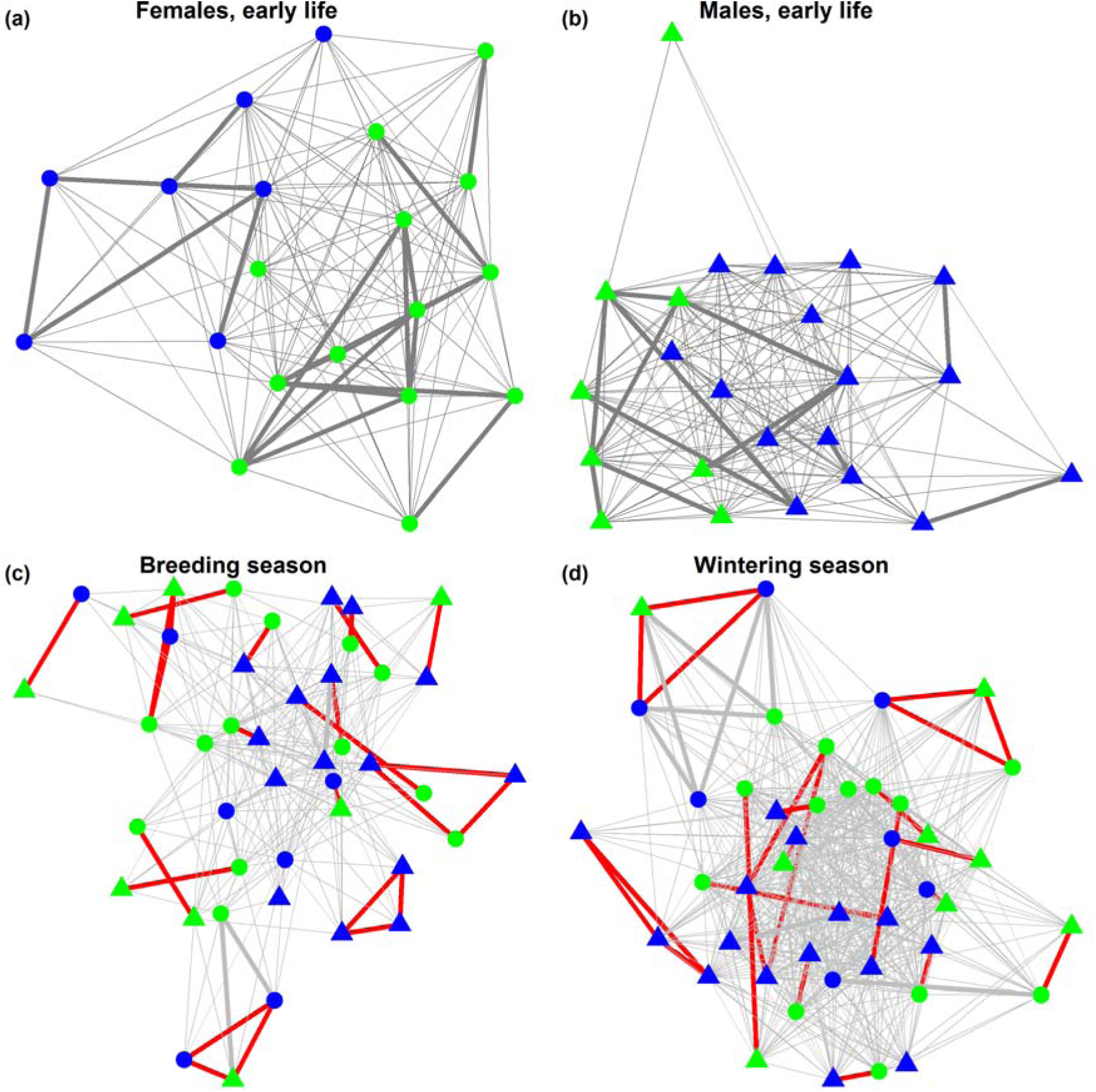
Nodes with the same colour indicate individuals coming from the same familiarity group. In the single-sex groups prior to pair formation, geese associated more with individuals from their own familiarity group as shown by the strong segregation of colours (and as reported in Kurvers et al., 2013). This pattern is, however, not present anymore in the breeding and wintering season. Circles represent females, and triangles males.

**Figure A3.**
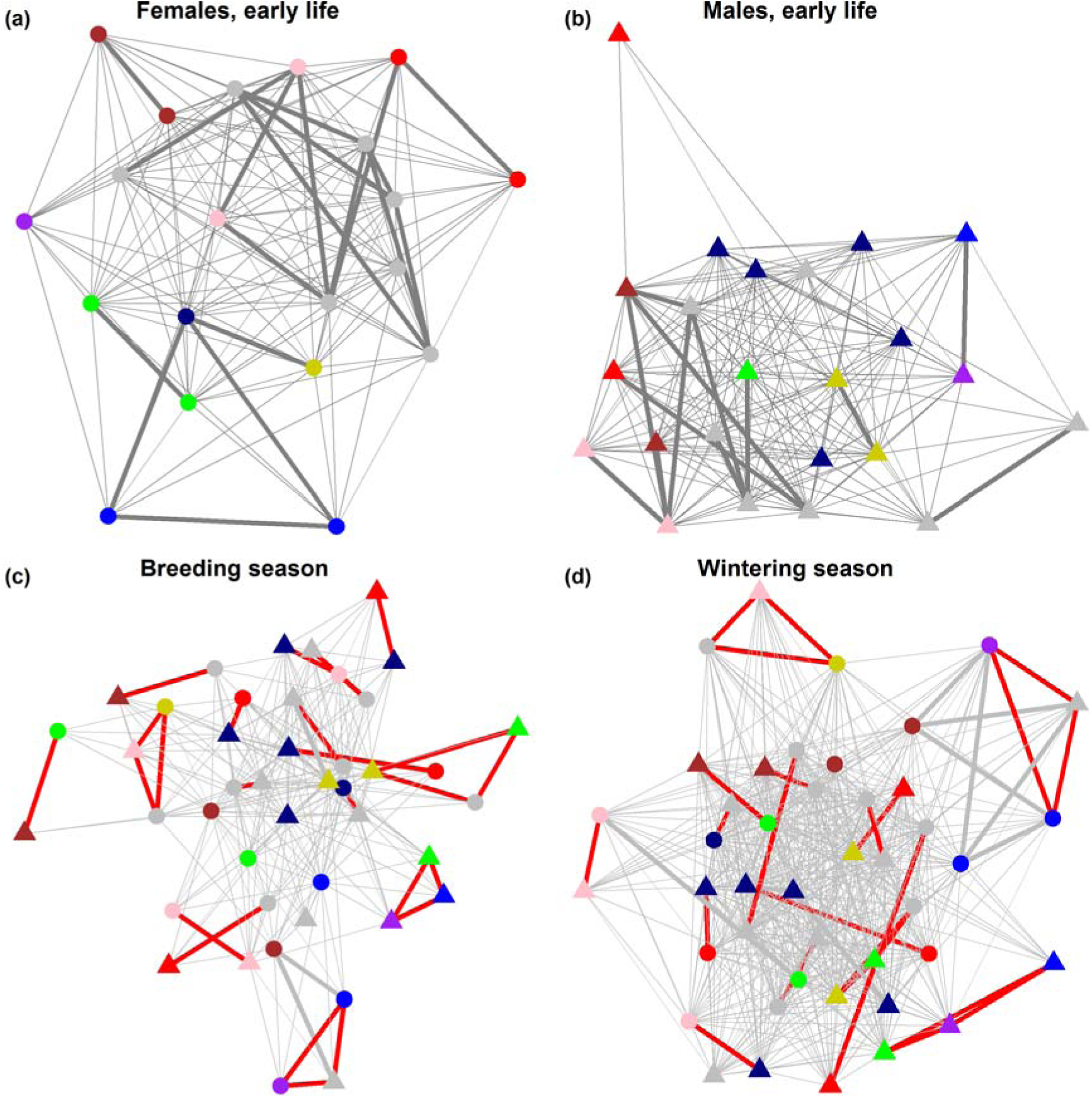
Nodes with the same colour indicate individuals with high genetic relatedness. Each cluster of colours has a mean genetic relatedness above 0.3. Grey-coloured nodes represent individuals not sharing a high genetic relatedness with any individual in the population. In the single-sex groups before pair formation, geese associated more with genetically related individuals colours (as reported in Kurvers et al., 2013). This pattern is, however, not present anymore in the breeding and wintering season. Circles represent females, and triangles males.

**Figure A4.**
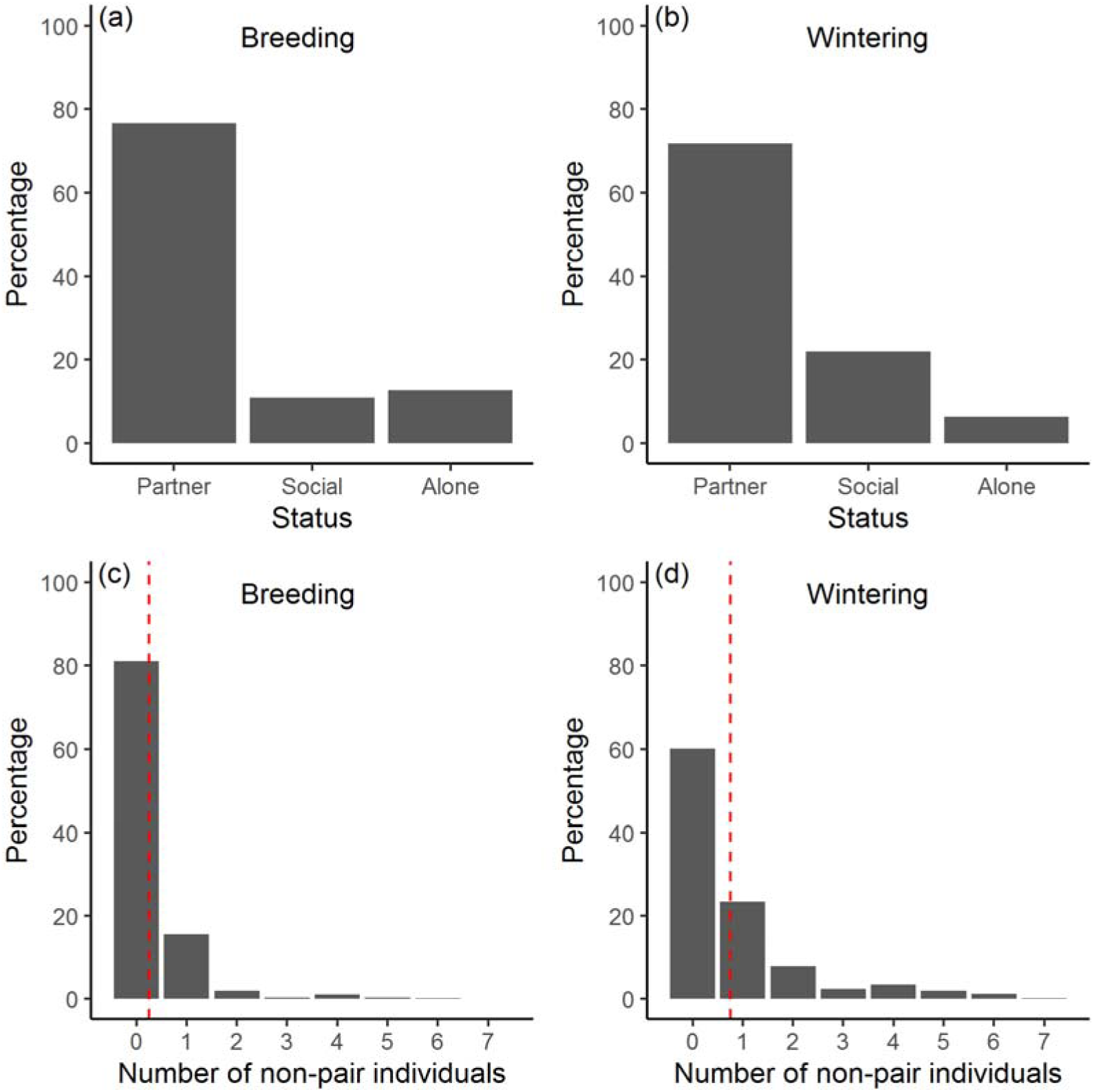
(a, b) The observed likelihood that when a paired individual was present on a patch that its partner was also present (‘partner’), that it was together with another individual but not its partner (‘social’), or that it was alone (‘alone’), for the (a) breeding, and (b) wintering season. (c, d) The observed likelihood of the number of individuals not belonging to the pair, which were present at a patch in the presence of a pair during the (c) breeding, and (d) wintering season. To illustrate, in the breeding season, in 80% of cases when pair members were observed together on a patch, there would be no other individuals (i.e., not belonging to that pair) present. In the wintering season, this percentage was around 60%. Red dashed lines represent the average number of non-paired geese present with a pair (mean during breeding: 0.26, during wintering: 0.76).

## References

Alexander, R. D. (1974). The Evolution of Social Behavior. Annual Review of Ecology and Systematics, 5(1), 325–383. doi: 10.1146/annurev.es.05.110174.001545

Anderholm, S., Marshall, R. C., van der Jeugd, H. P., Waldeck, P., Larsson, K., & Andersson, M. (2009a). Nest parasitism in the barnacle goose: evidence from protein fingerprinting and microsatellites. Animal Behaviour, 78(1), 167–174. doi: 10.1016/j.anbehav.2009.04.011

Anderholm, S., Waldeck, P., Van der Jeugd, H. P., Marshall, R. C., Larsson, K., & Andersson, M. (2009b). Colony kin structure and host-parasite relatedness in the barnacle goose. Molecular Ecology, 18(23), 4955–4963. doi: 10.1111/j.1365-294X.2009.04397.x

Andersson, M., Åhlund, M., & Waldeck, P. (2019). Brood parasitism, relatedness and sociality: a kinship role in female reproductive tactics. Biological Reviews, 94(1), 307–327.

Bejder, L., Fletcher, D., & Brager, S. (1998). A method for testing association patterns of social animals. Animal Behaviour, 56, 719–725.

Beletsky, L. D., & Orians, G. H. (1989). Familiar neighbors enhance breeding success in birds. Proceedings of the National Academy of Sciences, 86(20), 7933–7936. doi: 10.1073/pnas.86.20.7933

Black, J. M. (2001). Fitness consequences of long-term pair bonds in barnacle geese: monogamy in the extreme. Behavioral Ecology, 12(5), 640–645. doi: 10.1093/beheco/12.5.640

Black, J. M., Choudhury, S., & Owen, M. (1996). Do barnacle geese benefit from lifelong monogamy? Oxford Ornithology Series, 6, 91–117.

Black, J. M., & Owen, M. (1989). Parent Offspring Relationships in Wintering Barnacle Geese. Animal Behaviour, 37, 187–198. doi: 10.1016/0003-3472(89)90109-7

Black, J. M., & Owen, M. (1995). Reproductive performance and assortative pairing in relation to age in Barnacle geese. Journal of Animal Ecology, 64(2), 234–244. doi: 10.2307/5758

Black, J. M., Prop, J., & Larsson, K. (2014). The barnacle goose. Bloomsbury, UK: Bloomsbury Publishing.

Blumstein, D. T., Wey, T. W., & Tang, K. (2009). A test of the social cohesion hypothesis: interactive female marmots remain at home. Proceedings of the Royal Society B-Biological Sciences, 276(1669), 3007–3012. doi: 10.1098/rspb.2009.0703

Cameron, E. Z., Setsaas, T. H., & Linklater, W. L. (2009). Social bonds between unrelated females increase reproductive success in feral horses. Proceedings of the National Academy of Sciences of the United States of America, 106(33), 13850–13853. doi: 10.1073/pnas.0900639106

Carter, A. J., Macdonald, S. L., Thomson, V. A., & Goldizen, A. W. (2009). Structured association patterns and their energetic benefits in female eastern grey kangaroos, *Macropus giganteus*. Animal Behaviour, 77(4), 839–846.

Carter, K. D., Brand, R., Carter, J. K., Shorrocks, B., & Goldizen, A. W. (2013). Social networks, long-term associations and age-related sociability of wild giraffes. Animal Behaviour, 86(5), 901–910. doi: https://doi.org/10.1016/j.anbehav.2013.08.002

Choudhury, S., & Black, J. M. (1993). Mate selection behavior and sampling strategies in geese. Animal Behaviour, 46(4), 747–757.

Choudhury, S., & Black, J. M. (1994). Barnacle geese preferentially pair with familiar associates from early-life. Animal Behaviour, 48(1), 81–88.

Choudhury, S., Jones, C. S., Black, J. M., & Prop, J. (1993). Adoption of young and intraspecific nest parasitism in barnacle geese. The Condor, 95(4), 860–868.

Chuang, M.-F., Kam, Y.-C., & Bee, M. A. (2017). Territorial olive frogs display lower aggression towards neighbours than strangers based on individual vocal signatures. Animal Behaviour, 123, 217–228.

Connor, R. C., Cioffi, W. R., Randic, S., Allen, S. J., Watson-Capps, J., & Krutzen, M. (2017). Male alliance behaviour and mating access varies with habitat in a dolphin social network. Scientific Reports, 7, 46354. doi: 10.1038/srep46354

Croft, D. P., James, R., & Krause, J. (2008). Exploring animal social networks. Princeton, New Jersey: Princeton University Press.

Drent, R. H., & Prop, J. (2008). Barnacle goose *Branta leucopsis* survey on Nordenskiöldkysten, west Spitsbergen 1975–2007: breeding in relation to carrying capacity and predator impact. Circumpolar Stud, 4, 59–83.

Drent, R. H., & Swierstra, P. (1977). Goose flocks and food finding: field experiments with Barnacle Geese in winter. Wildfowl, 28, 15–20.

Farine, D. R. (2013). Animal social network inference and permutations for ecologists in R using asnipe. Methods in Ecology and Evolution, 4(12), 1187–1194.

Farine, D. R. (2017). A guide to null models for animal social network analysis. Methods in Ecology and Evolution, 8(10), 1309–1320. doi: 10.1111/2041-210x.12772

Farine, D. R., & Whitehead, H. (2015). Constructing, conducting and interpreting animal social network analysis. Journal of Animal Ecology, 84(5), 1144–1163.

Firth, J. A., & Sheldon, B. C. (2016). Social carry-over effects underpin trans-seasonally linked structure in a wild bird population. Ecology letters, 19(11), 1324–1332.

Forslund, P. (1993). Vigilance in relation to brood size and predator abundance in the barnacle goose, *Branta leucopsis*. Animal Behaviour, 45(5), 965–973.

Forslund, P., & Larsson, K. (1995). Intraspecific nest parasitism in the barnacle goose: behavioural tactics of parasites and hosts. Animal Behaviour, 50(2), 509–517.

Franks, D. W., Ruxton, G. D., & James, R. (2010). Sampling animal association networks with the gambit of the group. Behavioral Ecology and Sociobiology, 64(3), 493–503. doi: 10.1007/s00265-009-0865-8

Frigerio, D., Weiss, B., & Kotrschal, K. (2001). Spatial proximity among adult siblings in greylag geese (*Anser anser*): evidence for female bonding? Acta ethologica, 121–125.

Gilby, I. C., Brent, L. J. N., Wroblewski, E. E., Rudicell, R. S., Hahn, B. H., Goodall, J., & Pusey, A. E. (2013). Fitness benefits of coalitionary aggression in male chimpanzees. Behavioral Ecology and Sociobiology, 67(3), 373–381. doi: 10.1007/s00265-012-1457-6

Ginsberg, J. R., & Young, T. P. (1992). Measuring association between individuals or groups in behavioral studies. Animal Behaviour, 44(2), 377–379. doi: 10.1016/0003-3472(92)90042-8

Grabowska-Zhang, A., Sheldon, B., & Hinde, C. (2012). Long-term familiarity promotes joining in neighbour nest defence. Biology Letters, 8(4), 544–546.

Griffiths, S. W., Brockmark, S., Höjesjö, J., & Johnsson, J. (2004). Coping with divided attention: the advantage of familiarity. Proceedings of the Royal Society of London. Series B: Biological Sciences, 271(1540), 695–699.

Henzi, S. P., Lusseau, D., Weingrill, T., van Schaik, C. P., & Barrett, L. (2009). Cyclicity in the structure of female baboon social networks. Behavioral Ecology and Sociobiology, 63(7), 1015–1021. doi: 10.1007/s00265-009-0720-y

Hoppitt, W. J., & Farine, D. R. (2018). Association indices for quantifying social relationships: how to deal with missing observations of individuals or groups. Animal Behaviour, 136, 227–238.

Ilany, A., Booms, A. S., & Holekamp, K. E. (2015). Topological effects of network structure on long-term social network dynamics in a wild mammal. Ecology letters, 18(7), 687–695.

Jaeger, R. G. (1981). Dear enemy recognition and the costs of aggression between salamanders. The American Naturalist, 117(6), 962–974.

Jonker, R. M., Zhang, Q., Van Hooft, P., Loonen, M. J., Van der Jeugd, H. P., Crooijmans, R. P., … Kraus, R. H. (2012). The development of a genome wide SNP set for the Barnacle goose *Branta leucopsis*. PLoS One, 7(7), e38412. doi: 10.1371/journal.pone.0038412

Kappeler, P. M. (2019). A framework for studying social complexity. Behavioral Ecology and Sociobiology, 73(1), 13. doi: 10.1007/s00265-018-2601-8

Kappeler, P. M., Barrett, L., Blumstein, D. T., & Clutton-Brock, T. H. (2013). Constraints and flexibility in mammalian social behaviour: introduction and synthesis. Philosophical Transactions of the Royal Society B: Biological Sciences, 368(1618), 20120337. doi: 10.1098/rstb.2012.0337

Kerth, G., Perony, N., & Schweitzer, F. (2011). Bats are able to maintain long-term social relationships despite the high fission-fusion dynamics of their groups. Proceedings of the Royal Society B, 278(1719), 2761–2767. doi: 10.1098/rspb.2010.2718

Kohn, G. M. (2017). Friends give benefits: autumn social familiarity preferences predict reproductive output. Animal behaviour, 132, 201–208.

Krackhardt, D. (1988). Predicting with networks: Nonparametric multiple regression analysis of dyadic data. Social Networks, 10, 359–381.

Kraus, R. H. S., Kerstens, H., Van Hooft, P., Crooijmans, R., Van Der Poel, J., Elmberg, J., … Groenen, M. (2011). Genome wide SNP discovery, analysis and evaluation in mallard (*Anas platyrhynchos*). BMC Genomics, 12(1), 150.

Krause, J., & Ruxton, G. (2002). Living in Groups. Oxford: Oxford University Press.

Kurvers, R. H., Straates, K., Ydenberg, R. C., van Wieren, S. E., Swierstra, P. S., & Prins, H. H. (2014). Social information use by barnacle geese *Branta leucopsis*, an experiment revisited. Ardea, 102(2), 173–181.

Kurvers, R. H. J. M., Adamczyk, V. M. A. P., Kraus, R. H. S., Hoffman, J. I., van Wieren, S. E., van der Jeugd, H. P., … Jonker, R. M. (2013). Contrasting context dependence of familiarity and kinship in animal social networks. Animal Behaviour, 86(5), 993–1001. doi: 10.1016/j.anbehav.2013.09.001

Kurvers, R. H. J. M., Eijkelenkamp, B., van Oers, K., van Lith, B., van Wieren, S. E., Ydenberg, R. C., & Prins, H. H. T. (2009). Personality differences explain leadership in barnacle geese. Animal Behaviour, 78(2), 447–453.

Kurvers, R. H. J. M., Prins, H. H. T., van Wieren, S. E., van Oers, K., Nolet, B. A., & Ydenberg, R. C. (2010). The effect of personality on social foraging: shy barnacle geese scrounge more. Proceedings of the Royal Society B-Biological Sciences, 277(1681), 601–608.

Kutsukake, N. (2006). The context and quality of social relationships affect vigilance behaviour in wild chimpanzees. Ethology, 112(6), 581–591.

Langenhof, M. R., & Komdeur, J. (2018). Why and how the early-life environment affects development of coping behaviours. Behavioral Ecology and Sociobiology, 72(3), 34. doi: 10.1007/s00265-018-2452-3

Leris, I., & Reader, S. M. (2016). Age and early social environment influence guppy social learning propensities. Animal Behaviour, 120, 11–19. doi: https://doi.org/10.1016/j.anbehav.2016.07.012

Lindström, J. (1999). Early development and fitness in birds and mammals. Trends in Ecology & Evolution, 14(9), 343–348. doi: https://doi.org/10.1016/S0169-5347(99)01639-0

Linklater, W. L., & Cameron, E. Z. (2009). Social dispersal but with philopatry reveals incest avoidance in a polygynous ungulate. Animal Behaviour, 77(5), 1085–1093.

Maher, C. R., & Burger, J. R. (2011). Intraspecific variation in space use, group size, and mating systems of caviomorph rodents. Journal of Mammalogy, 92(1), 54–64.

Milligan, B. G. (2003). Maximum-likelihood estimation of relatedness. Genetics, 163(3), 1153–1167.

Mitani, J. C. (2009). Male chimpanzees form enduring and equitable social bonds. Animal Behaviour, 77(3), 633–640. doi: https://doi.org/10.1016/j.anbehav.2008.11.021

Oksanen, J., Blanchet, G., Friendly, M., Kindt, R., Legendre, P., McGlinn, D., … Wagner, H. (2019). vegan: Community Ecology Package. R package version 2.5-4.

Olendorf, R., Getty, T., & Scribner, K. (2004). Cooperative nest defence in red–winged blackbirds: reciprocal altruism, kinship or by–product mutualism? Proceedings of the Royal Society of London. Series B: Biological Sciences, 271(1535), 177–182.

Owen, M., Black, J. M., & Liber, H. (1988). Pair bond duration and timing of its formation in barnacle geese (*Branta leucopsis*). In M. W. Weller (Ed.), Waterfowl in Winter. Minneapolis, Minnesota: University of Minnesota Press.

Owen, M., & Wells, R. (1979). Territorial behaviour in breeding geese--a re-examination of Ryder’s hypothesis. Wildfowl, 30(30), 20–26.

Percival, S. M. (1991). The Population-Structure of Greenland Barnacle Geese *Branta-Leucopsis* on the Wintering Grounds on Islay. Ibis, 133(4), 357–364.

Raveling, D. G., Sedinger, J. S., & Johnson, D. S. (2000). Reproductive success and survival in relation to experience during the first two years in Canada geese. The Condor, 102(4), 941–945.

Robertson, G. J., & Cooke, F. (1999). Winter philopatry in migratory waterfowl. The Auk, 116(1), 20–34.

Sachser, N., Dürschlag, M., & Hirzel, D. (1998). SOCIAL RELATIONSHIPS AND THE MANAGEMENT OF STRESS. Psychoneuroendocrinology, 23(8), 891–904. doi: https://doi.org/10.1016/S0306-4530(98)00059-6

Samelius, G., & Alisauskas, R. T. (2006). Sex-biased costs in nest defence behaviours by lesser snow geese (*Chen caerulescens*): consequences of parental roles? Behavioral ecology and sociobiology, 59(6), 805–810.

Scheiber, I. B. R., Hohnstein, A., Kotrschal, K., & Weiß, B. M. (2011). Juvenile Greylag Geese (Anser anser) Discriminate between Individual Siblings. PLOS ONE, 6(8), e22853. doi: 10.1371/journal.pone.0022853

Scheiber, I. B. R., Kotrschal, K., & Weiss, B. M. (2009). Benefits of family reunions: Social support in secondary greylag goose families. Hormones and Behavior, 55(1), 133–138. doi: 10.1016/j.yhbeh.2008.09.006

Scheiber, I. B. R., Weiss, B. M., Frigerio, D., & Kotrschal, K. (2005). Active and passive social support in families of greylag geese (*Anser anser*). Behaviour, 142, 1535–1557. doi: 10.1163/156853905774831873

Schulke, O., Bhagavatula, J., Vigilant, L., & Ostner, J. (2010). Social Bonds Enhance Reproductive Success in Male Macaques. Current Biology, 20(24), 2207–2210. doi: 10.1016/j.cub.2010.10.058

Shizuka, D., Chaine, A. S., Anderson, J., Johnson, O., Laursen, I. M., & Lyon, B. E. (2014). Across-year social stability shapes network structure in wintering migrant sparrows. Ecology Letters, 17(8), 998–1007.

Silk, J. B. (2007). Social components of fitness in primate groups. Science, 317(5843), 1347–1351.

Silk, J. B., Alberts, S. C., & Altmann, J. (2003). Social bonds of female baboons enhance infant survival. Science, 302(5648), 1231–1234. doi: 10.1126/science.1088580

Silk, J. B., Beehner, J. C., Bergman, T. J., Crockford, C., Engh, A. L., Moscovice, L. R., … Cheney, D. L. (2009). The benefits of social capital: close social bonds among female baboons enhance offspring survival. Proceedings of the Royal Society B-Biological Sciences, 276(1670), 3099–3104. doi: 10.1098/rspb.2009.0681

Silk, J. B., Beehner, J. C., Bergman, T. J., Crockford, C., Engh, A. L., Moscovice, L. R., … Cheney, D. L. (2010). Strong and Consistent Social Bonds Enhance the Longevity of Female Baboons. Current Biology, 20(15), 1359–1361. doi: 10.1016/j.cub.2010.05.067

Siracusa, E. R., Wilson, D. R., Studd, E. K., Boutin, S., Humphries, M. M., Dantzer, B., … McAdam, A. G. (2019). North American red squirrels mitigate costs of territory defence through social plasticity. Animal behaviour, 151, 29–42.

Stanley, C. R., Mettke-Hofmann, C., Hager, R., & Shultz, S. (2018). Social stability in semiferal ponies: networks show interannual stability alongside seasonal flexibility. Animal Behaviour, 136, 175–184.

Stanton, M. A., & Mann, J. (2012). Early social networks predict survival in wild bottlenose dolphins. PloS one, 7(10), e47508.

Szipl, G., Depenau, M., Kotrschal, K., Hemetsberger, J., & Frigerio, D. (2019). Costs and benefits of social connectivity in juvenile Greylag geese. Scientific Reports, 9(1), 12839. doi: 10.1038/s41598-019-49293-9

Temeles, E. J. (1994). The role of neighbours in territorial systems: when are they ‘dear enemies’? Animal Behaviour, 47(2), 339–350.

van der Jeugd, H. P. (2001). Large barnacle goose males can overcome the social costs of natal dispersal. Behavioral Ecology, 12(3), 275–282.

van der Jeugd, H. P., & Blaakmeer, K. B. (2001). Teenage love: the importance of trial liaisons, subadult plumage and early pairing in barnacle geese. Animal Behaviour, 62, 1075–1083.

van der Jeugd, H. P., van der Veen, I. T., & Larsson, K. (2002). Kin clustering in barnacle geese: familiarity or phenotype matching? Behavioral Ecology, 13(6), 786–790. doi: 10.1093/beheco/13.6.786

Wang, J. L. (2011). COANCESTRY: a program for simulating, estimating and analysing relatedness and inbreeding coefficients. Molecular Ecology Resources, 11(1), 141–145. doi: 10.1111/j.1755-0998.2010.02885.x

Whitehead, H. (2008). Analyzing animal societies: Quantative methods for vertebrate social analysis. Chicago, Illinois: University of Chicago Press.

Ydenberg, R. C., Giraldeau, L. A., & Falls, J. B. (1988). Neighbours, strangers, and the asymmetric war of attrition. Animal Behaviour, 36(2), 343–347. doi: https://doi.org/10.1016/S0003-3472(88)80004-6

Zeus, V. M., Reusch, C., & Kerth, G. (2018). Long-term roosting data reveal a unimodular social network in large fission-fusion society of the colony-living Natterer’s bat (*Myotis nattereri*). Behavioral Ecology and Sociobiology, 72(6), 99. doi: 10.1007/s00265-018-2516-4

